# Prior exposure to microcystin alters host gut resistome and is associated with dysregulated immune homeostasis in translatable mouse models

**DOI:** 10.1101/2021.12.17.472989

**Authors:** Punnag Saha, Dipro Bose, Vitalii Stebliankin, Trevor Cickovski, Ratanesh K. Seth, Dwayne E. Porter, Bryan W. Brooks, Kalai Mathee, Giri Narasimhan, Rita Colwell, Geoff I. Scott, Saurabh Chatterjee

## Abstract

The increased propensity of harmful algal blooms (HABs) and exposure from HABs-cyanotoxin causes human toxicity. It has been associated with the progression of several diseases that encompass the liver, kidneys, and immune system. Recently, a strong association of cyano-HAB toxicity with the altered host gut microbiome has been shown. We tested the hypothesis that prior exposure to cyanotoxin microcystin may alter the microbiome and induce microbiome-host-resistome crosstalk. Using both wild-type and humanized mice, we show that the mice exposed to microcystin had an altered microbiome signature that harbored antimicrobial resistance genes. Host resistome phenotypes such as *mefA, msrD, mel, ant6*, and *tet40* increased in diversity and relative abundance following microcystin exposure. Interestingly, the increased abundance of these genes was traced to resistance to common antibiotics such as tetracycline, macrolides, glycopeptide, and aminoglycosides, crucial for modern-day treatment for several diseases. Increased abundance of these genes was positively associated with increased expression of PD1, a T-cell homeostasis marker, and pleiotropic inflammatory cytokine IL-6 with a concomitant negative association with immunosurveillance markers IL7 and TLR2. Microcystin exposure also caused decreased TLR2, TLR4, and REG3G expressions, increased immunosenescence, and higher systemic levels of IL-6 in both wild-type and humanized mice. In conclusion, the results show a first-ever characterization of the host resistome of microcystin exposure and its connection to host immune status and antibiotic resistance. The results may be crucial for understanding the ability of exposed subjects to fight future bacterial infections and the progression of the debilitating disease in hospital settings.

## 1 Introduction

Occurrence of harmful cyanobacterial algal blooms (HABs) has been increasingly found over the past decades across the world^1^. In the United States, the presence of cyanobacterial HABs has been reported in numerous inland lakes and rivers^2–4^. Anthropogenic activities include the release of wastewater effluents, agricultural runoffs containing fertilizers and pesticides, and urban runoffs into water bodies, resulting in increased eutrophication. This eutrophication in combination with climate change-associated global warming extensively favors the excessive proliferation of cyanobacteria, which ultimately leads to bloom formation^5–7^. HABs are well-known producers of diverse types of cyanotoxins in the water systems as secondary metabolites. These cyanotoxins are often broadly categorized according to their mode of toxicity, e.g., hepatotoxins (microcystin, cylindrospermopsin, nodularin), neurotoxins (anatoxin-a, saxitoxin), dermatoxins (aplysiatoxin), etc.,^8^. Possible routes of potential cyanotoxin exposure in humans include consuming contaminated drinking water, ingestion or inhalation, and dermal contact with these toxins for recreational purposes, e.g., swimming^9^. However, drinking contaminated water remains the plausible major route of exposure for freshwater cyanotoxins. As mentioned by CDC, exposure to cyanotoxins can lead to multiple adverse effects in humans, including gastroenteritis, nausea, vomiting, respiratory distress, dermatitis, headache, and liver problems depending on the exposure route and cyanotoxin itself. Also, infants and children are more likely to be at higher risk of cyanotoxin-associated pathological outcomes than adults as they consume more water per body weight^10^.

Among the wide array of cyanotoxins released by HABs, microcystin is the most prevalent and well-studied toxin, whereas microcystin-LR (MC) is regarded as the most toxic variant among the 240 reported congeners of microcystin^9,11,12^. MC is primarily produced by the cyanobacterial species *Microcystis aeruginosa*, although many other genera including *Anabaena, Nostoc, Planktothrix, Hapalosiphon*, and *Phormidium* are also reported as MC producers^13^. Being a water-soluble cyclic heptapeptide, MC uptake is carried out by organic anion transport polypeptides (OATPs)-dependent mechanism in various organ systems e.g., liver, kidneys, and intestines as well as the brain^14–16^. Mechanistically, MC inhibits the catalytic activity of major cellular dephosphorylating enzymes Ser/Thr protein phosphatase 1 (PP1) and protein phosphatase 2A (PP2A) by covalent modifications^17–19^. Blockage of PP1 and PP2A by MC hampers the cell cycle regulation, proliferation, cytoskeletal assembly, and a plethora of downstream signaling pathways^20,21^. Over the past few years, our research group has worked extensively on novel mechanistic insights of MC-associated toxicity using both *in vivo* and *in vitro* models. Our findings show the role of MC-mediated hepatotoxicity in both adult^22^ and juvenile mice^23^, renal toxicity^24^, gastrointestinal toxicity^25,26^, and neurotoxicity^27^, leading to progression of Nonalcoholic fatty liver disease (NAFLD) to an advanced and more progressive stage of pathology.

Previous studies by our research group and several others have reported that exposure to MC significantly altered gut bacteriome in experimental murine models^25,28,29^. Gut dysbiosis affects the host health by exacerbating the pathological outcomes in gastrointestinal disorders, liver diseases, and metabolic conditions like obesity and diabetes and increases the risk of anti-microbial resistance, which has been a global health concern ^30,31^.

Recent reports have identified that along with antibiotics, various other environmental exposures like pesticides, insecticides, and heavy metals also influence the increase in anti-microbial resistance genes or ARGs^32–37^. The sustained exposure to these factors is known to create a selection pressure on the environmental and gut bacteria, allowing them to express these ARGs for survival^38^. The expressed ARGs may be resistant against a single antibiotic or encode multiple drug resistance and can be transferred between bacteria using mobile genetic elements (MGE) by horizontal gene transfer (HGT) persisting for a prolonged period^39,40^. Increased emergence of antibiotic-resistant bacteria in clinics, hospitals, and community environments leads to delayed and ineffective treatment, often leading to mortality^41,42^.

Effects of environmental toxins like MC on gut dysbiosis have been widely studied. However, its effect on gut resistome remains unknown. In addition, the association between increased ARGs and immunological markers in MC exposed condition has not been studied earlier. This led us to hypothesize that MC exposure might lead to increased anti-microbial resistance in the gut, and any future bacterial infection in persons pre-exposed to MC might pose a serious challenge in terms of treatment, which immensely emphasizes the clinical importance of the present study. In this study, we investigated the effect of early life MC exposure on modulating gut bacteriome and resistome and its association with intestinal pathology and systemic inflammation using an experimental murine model. We also used a novel focus area in studying the immune-phenotypical changes following MC exposure in humanized mice to ascertain the translatability of MC exposure in humans.

## 2. Materials and methods

### 2.1. Materials

Microcystin-LR (MC) used in this study was purchased from Cayman Chemical Company (Ann Arbor, MI, USA).

### 2.2. Animals

Pathogen-free, male, juvenile, wild-type (WT) C57BL/6J mice and female, adult, NSG™ mice (engrafted with human CD34^+^ hematopoietic stem cells) were purchased from Jackson Laboratories (Ban Harbor, ME, USA). All mice experiments were conducted by strictly following the National Institutes of Health (NIH) guidelines for humane care and use of laboratory animals and local Institutional Animal Care and Use Committee (IACUC) standards. The animal handling procedures for this study were approved by the University of South Carolina (Columbia, SC, United States). Upon arrival, all mice were housed in a 22–24°C temperature-controlled room with a 12 h light/12 h dark cycle and had ad libitum access to food and water. Upon completion of dosing, all mice were sacrificed. The distal parts of the small intestine were collected from each sacrificed mouse and immediately fixed in 10% neutral buffered formaldehyde (Sigma-Aldrich, St. Louis, MO, USA). Also, serum samples were prepared from freshly collected blood and were kept at −80°C. Fecal pellets were collected from the colon of each mouse, snap-frozen immediately, and preserved at −80 °C for microbiome analysis.

### 2.3. Experimental murine model of MC exposure

Upon arrival, all mice were acclimatized for one week. For this study, 12 WT mice (4 weeks old) and 8 humanized NSG™ mice (18 weeks old) mice were randomly divided into two groups (n=6; n=4 respectively). For both WT and humanized NSG™ mice, the control group of mice (CONTROL; Hu-CONTROL respectively) received only vehicle [Phosphate buffered saline (PBS)] whereas the treated group (MC; Hu-MC respectively) of mice were dosed with microcystin-LR (7μg/ kg body weight; dissolved in ethanol and then diluted in PBS) for a continuous 2 weeks by oral gavage route. Post MC treatment, all WT mice were rested for 4 weeks to grow till adulthood and then euthanized at the age of 10 weeks. However, all humanized NSG™ mice were euthanized immediately after MC treatment was completed. Both WT and humanized NSG™ mice were fed with only the Chow diet throughout the study.

#### 2.3.1. Rationale for using humanized NSG^TM^ mice transplanted with human CD34^+^ cells

MC exposure in WT mice, the associated microbiome, and resistome changes may not reflect the true human immune phenotype changes. To ascertain such translatable effects in humans following MC exposure, we used humanized NSG^TM^ mice with a known immune phenotype that reflects the humanized T and B cells^43^. The difference in the age of mice (WT vs the humanized NSG^TM^ mice) did not influence the results since we solely aimed to look into the translatability of the immune phenotype (T cell homeostasis/immunosenescence, CD28, CD57); (inflammation-Systemic IL-6) following MC exposure and not focus on the microbiome-led changes.

### 2.4. Bacteriome analysis

As mentioned in one of our previous works, raw reads were generated by the vendor CosmosID Inc. (Germantown, MD, USA) using the fecal pellets obtained from the WT experimental mice only^44^. Bacteriome analysis was then completed with the help of the MetaWRAP pipeline^45^. Raw reads were first trimmed using Trim Galore^46^ (version 0.6.7, default parameters). Host reads were eliminated with BMTagger 1.1.0, with the help of the Mouse Genome Database^47^. MegaHit 1.2.9^48^ was then used for genome assembly, followed by Kraken2^49^ which mapped these whole-genome sequences to the NCBI Bacteria Database^50^, producing a list of taxa and abundances.

### 2.5. ARG family detection and analysis

#### 2.5.1. Resistome profile

The anti-microbial resistance profile was consolidated using two methods. The first approach involves aligning metagenomic reads from each subject against 7,868 antibiotic-resistance genes from the anti-microbial database for high-throughput sequencing MEGARes version 2.0^51^. Metagenomic reads were mapped against the references using Bowtie2^52^, and the coverage of each gene in a sample was accessed with Samtools^53^. From the alignment files, we computed the reads per kilobase million (RPKM) for each gene in a sample and normalized the profile across samples to adapt to 1. A gene was considered a part of the profile only if the corresponding read coverage was at least 70%. The pipeline was executed using the PeTRi high-performance computing framework^54^.

The second method involves the CosmosID method. Raw metagenomic sequencing reads were trimmed using fastp, with fastp_qualified_quality of 15 and fastp_cut_mean_quality of 15. Trimmed reads were assembled using MEGAHIT using default parameters, and Metagenome-Assembled Genomes were generated through binning with MetaBAT2 using default parameters. The quality of MAGs was assessed using QUAST and BUSCO using default parameters for both. MAGs were run through CosmosID’s ARG pipeline for anti-microbial resistance identification, and the CosmosID-Hub Microbiome Platform for taxonomic identification. Output MAGs were then run through CosmosID’s ARG pipeline. For identification of ARGs and, the assembled genomes were screened against the Resfinder ARG database. ARGs were considered as present if their sequences matched with the assembled genome at >90% Nucleotide identity and >60% Alignment coverage of the gene’s sequence length. MAGs were directly analyzed by CosmosID-HUB Microbiome Platform (CosmosID Inc., Germantown, MD) described elsewhere^55,56^ for multi-kingdom microbiome analysis and profiling of antibiotic resistance and virulence genes and quantification of organisms’ relative abundance. Briefly, the system utilizes curated genome databases and a high-performance data-mining algorithm that rapidly disambiguates hundreds of millions of metagenomic sequence reads into the discrete microorganisms engendering the particular sequences.

The resistome profiles obtained by those two methods were concatenated and re-normalized. Shannon’s and Simpson’s diversity indices of resistome profiles were computed with the vegan R package^57^.

#### 2.5.2. Genes provenance

To identify the provenance of ARGs we performed de-novo assembly followed by metagenomic binning, genomes annotation, and blast search. First, metagenomic reads from each sample were assembled into contigs using the Megahit tool^48^. Second, we aggregated the contigs from every sample, and re-mapped the metagenomic reads against the assembled reference with minimap2^58^. Third, the alignment file of re-mapped reads was used as input to the Metabat2 adaptive binning tool to cluster contigs into draft genomes^59^. Next, each reconstructed genome was annotated with phylogenetic information using PhyloPhlAn 3.0^60^. Finally, we queried the ARGs against the assembled genomes using BLAST 2.12.0+. The microbial species were considered a likely source of the ABR gene if the e-value from a blast alignment against the corresponding assembled genome is less than 0.01.

#### 2.5.3. Confounder identification

Conditional independence tests were performed to determine if the correlation between the abundance of anti-microbial genes and immune markers was dictated by the microcystin exposure. In particular, we identified the variables that satisfy the causal Markov conditions, i.e., every variable is independent of its non-descendants conditional on parents^61^. The microcystin exposure (M) was considered a common cause (confounder) if the abundance of a resistance gene (G) is independent of immune marker (I) when condition on M but correlated otherwise. Such an approach is the first iteration of the causal structural learning algorithm^62^, which was successfully adapted for inferring causal relationships in microbiomes^63^. The algorithm was executed to only identify first-order dependencies due to the small size of the analyzed dataset. Conditional independence tests were performed with the blearn R package with Mutual Information (cond. Gauss.) method^64^.

### 2.6. Laboratory methods

#### 2.6.1. Quantitative real-time polymerase chain reaction (qRT-PCR)

Levels of gene expression in the small intestine tissue samples were measured by the two-step qRT-PCR protocol. Firstly, all tissue samples were homogenized in TRIzol reagent (Invitrogen, Rockford, IL, USA) and then centrifuged to eliminate any sort of exces tissue particles and debris. Following homogenization and centrifugation procedure,; total RNA from the individual sample was isolated and purified by using RNAse mini kit columns (Qiagen, Valencia, CA, USA) as per the manufacturer’s protocol. Then, purified RNA (1000 ng) was converted to cDNA by using the iScript cDNA synthesis kit (Bio-Rad, Hercules, CA, USA) following the manufacturer’s standard procedure. Finally, qRT-PCR was carried out with the gene-specific mouse primers using SsoAdvanced SYBR Green supermix (Bio-Rad, Hercules, CA, USA) and CFX96 thermal cycler (Bio-Rad, Hercules, CA, USA). Threshold cycle (Ct) values for the selected genes were normalized against 18S (internal control) values in the same sample. The relative fold change was calculated by the 2-ΔΔCt method using the vehicle-treated groups (CONTROL, and Hu-CONTROL) of mice as control. The sequences (5’-3’ orientation) for the mouse-specific and human-specific primers used for real-time PCR are mentioned in Table 1.

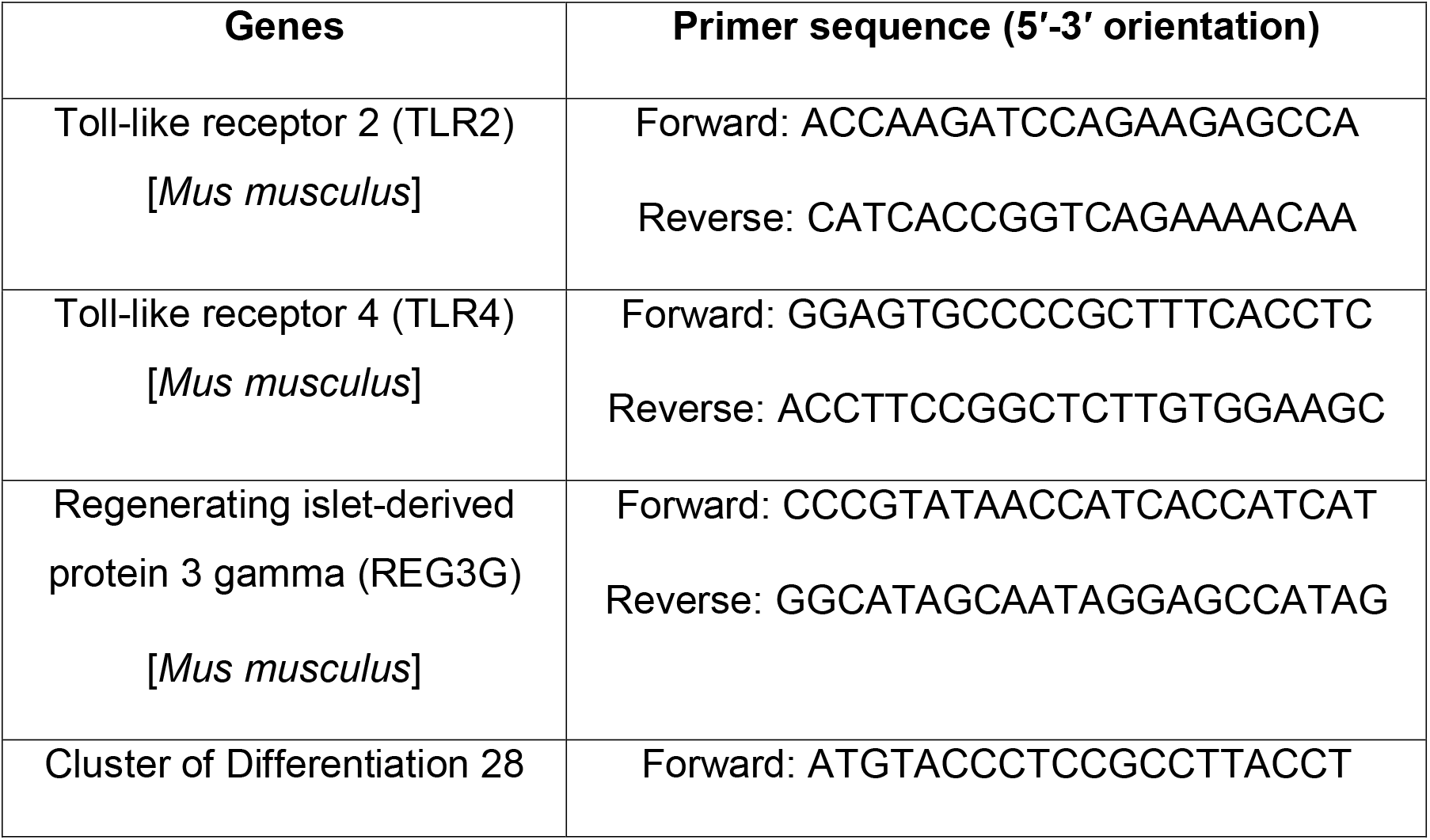
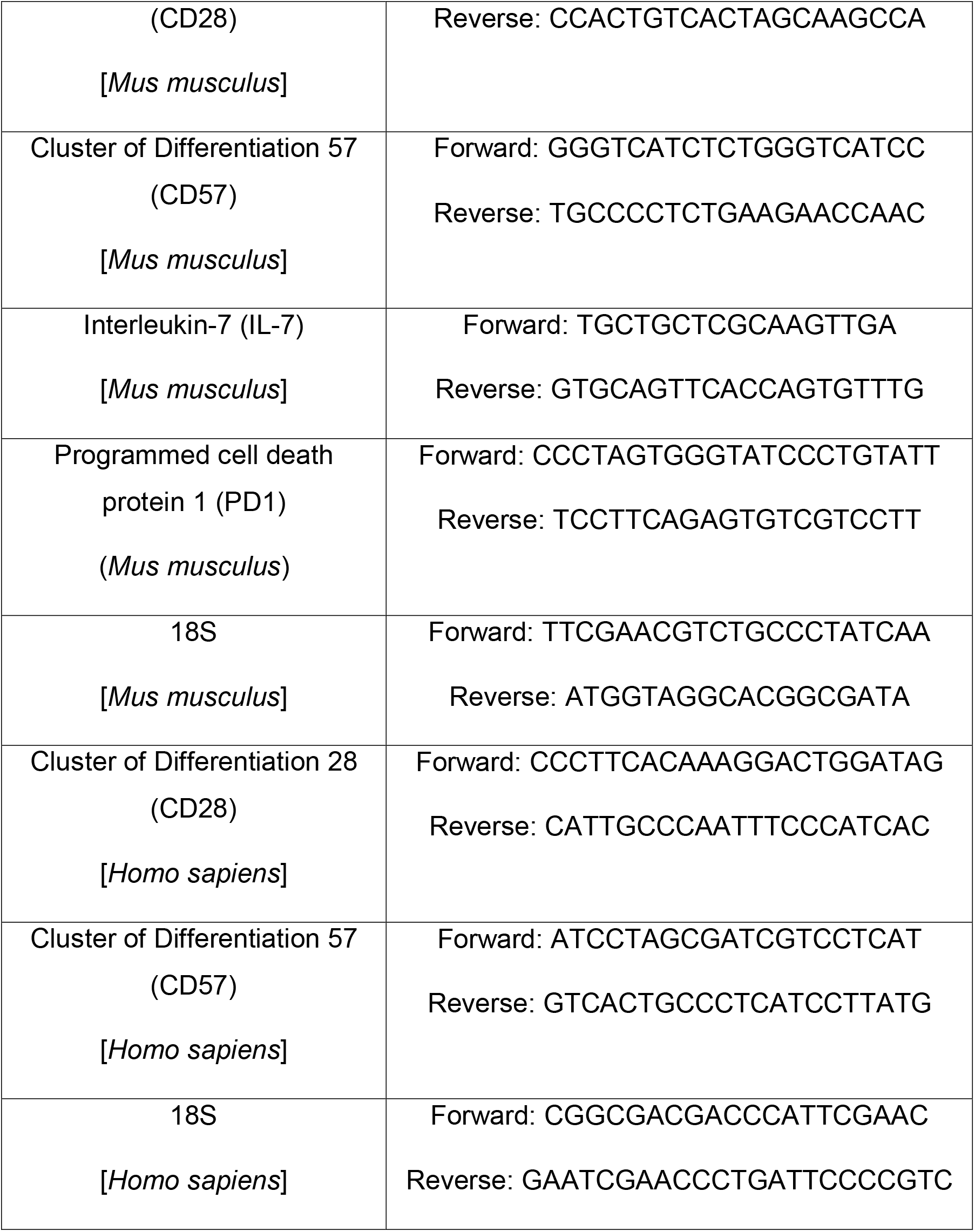

#### 2.6.2. Enzyme-Linked Immunosorbent Assay (ELISA)

For both the CONTROL and MC groups of mice, a mouse-specific ELISA Kit (Catalogue No. KE10007; ProteinTech, Rosemont, IL, USA) was used whereas for the Hu-CONTROL and Hu-MC groups of mice, a human-specific ELISA Kit (Catalogue No. KE00139; ProteinTech, Rosemont, IL, USA) was used to determine the serum IL-6 level following the manufacturer’s protocol.

### 2.7. Statistical analyses

All statistical analyses for this study were performed by using GraphPad Prism software (San Diego, CA, USA), and data were presented as mean ± SEM. For determining inter-group comparison, Unpaired t-test (two-tailed tests with equal variance) and one-way analysis of variance (ANOVA) were performed, followed by Bonferroni–Dunn post-hoc corrections analysis. Box and whisker plots of relative abundance pairwise comparisons were complemented with Wilcoxon rank-sum test p-values using *ggpubr* R package^65^. For all analyses, p ≤ 0.05 was considered statistically significant.

## 3. Results

### 3.1. Early life exposure to MC causes an altered gut bacteriome pattern in mice

First, we wanted to determine whether oral administration of MC in juvenile, WT mice for a continuous 2-week period resulted in any alteration of the gut bacteriome in adulthood. Next-Generation Sequencing was performed using the fecal pellets of mice to establish the detailed gut bacteriome profile for both experimental groups.

At the Phylum level, our results showed a distinctly different and altered relative abundance of gut commensals for both CONTROL and MC groups of mice (Figure 1. A.). We observed markedly decreased relative abundance of Bacteroidetes (Figure 1. B., p=0.39), Firmicutes (Figure 1. C., p=0.18) [statistically not significant], and Proteobacteria (Figure 1. D., p=0.0022) in the MC group compared to the CONTROL group. However relative abundance of both phyla Actinobacteria (Figure 1. D., p=0.0087) and Verrucomicrobia (Figure 1. F., p=0.015) were found to be significantly increased in the MC-exposed mice group when compared to the CONTROL group.

**Figure 1.**
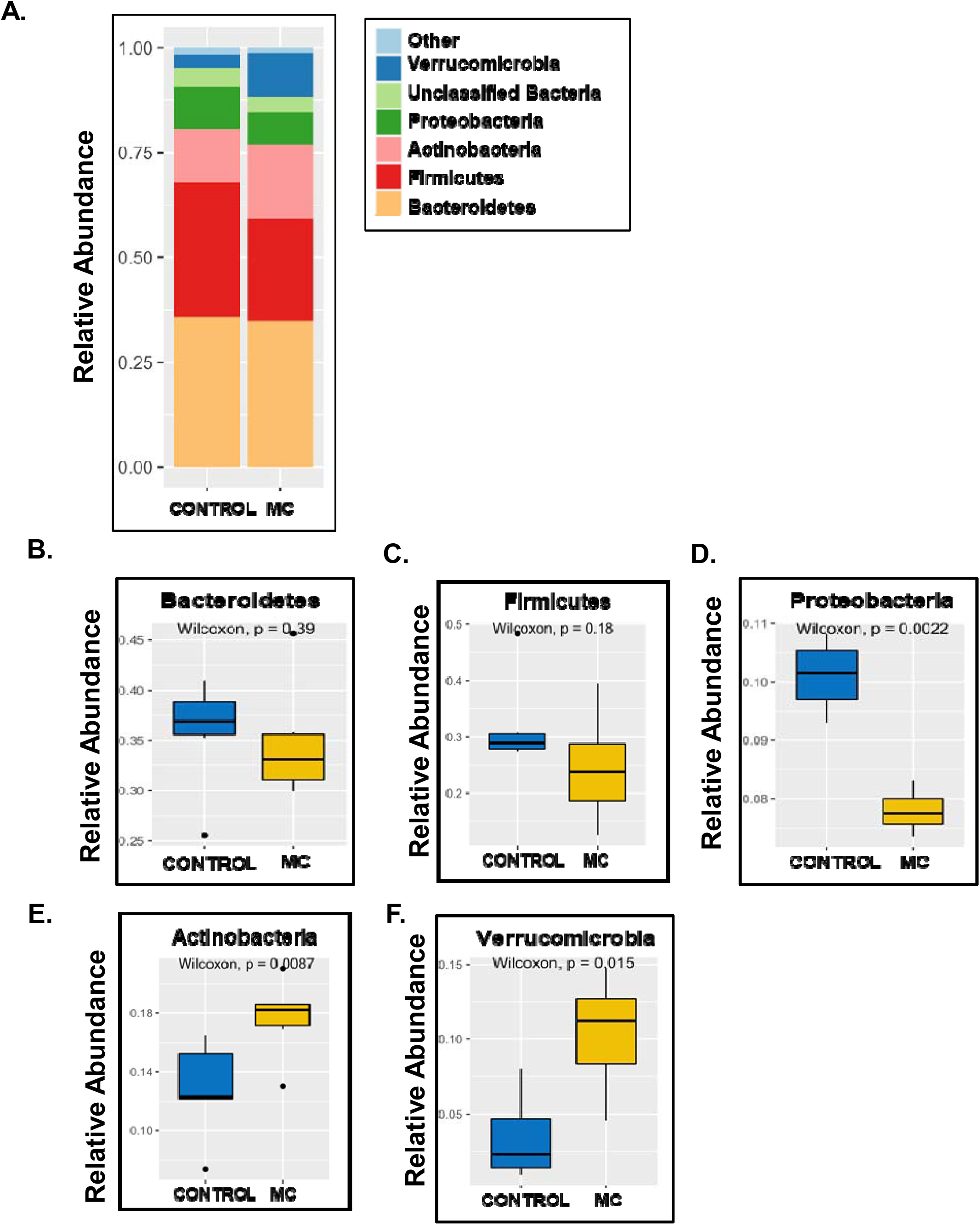
Relative abundance at the Phylum level. **(A.)** Bar plot showing relative abundance at the Phylum level for CONTROL (WT mice treated with vehicle only) and MC (WT mice orally administered with MC for 2 weeks) groups. Box and whisker plots showing the relative abundance of different phyla including **(B.)** Bacteriodetes, **(C.)** Firmicutes, **(D.)** Proteobacteria, **(E.)** Actinobacteria, and **(F.)** Verrumicrobia.

Next, we detected a prominently altered profile of relative abundance at the species level for both CONTROL and MC groups too (Figure 2. A.). *Akkermania muciniphilia* is an important member of gut microflora that primarily degrades the mucus layer of the intestinal lining. Under healthy conditions, this mucin degradation encourages the host’s intestinal Goblet cells to produce more mucin, which results in protecting the host from enteric pathogen adherence, thereby maintaining the gut barrier integrity and overall gut immune health^66^. However, our result showed a marked increase in the relative abundance of *A. muciniphilia* in the MC group compared to the CONTROL group (Figure 2. B., p=0.015). *Bacteroides thetaiotaomicron*, is a “friendly” commensal that particularly helps in carbohydrate metabolism in the host under normal physiological conditions^67^. However, under stressed conditions e.g., when the mucosal lining of the intestine is breached, *B. thetaiotaomicron* can also turn into an opportunistic pathogen leading to various infections, bacteremia in the host^67,68^, and most importantly this bacterium is associated with harboring ARGs in the gut^69^. In our study, we observed a significantly increased relative abundance of *B. thetaiotaomicron* in the MC group compared to the CONTROL group (Figure 2. C., p=0.0043). *Lactobacillus johnsonii* is a well-known “psychobiotic” bacterium which not only improves gut immune health but also exerts psychoactive effects as shown in a recent study^70^. Our results showed a markedly decreased relative abundance of *L. johnsonii* in the MC-exposed mice compared to the CONTROL mice (Figure 2. E., p=0.026). *Lactococcus lactis*, another prominent gut commensal with anti-inflammatory roles in the gut^71^, was also found to be significantly decreased in relative abundance in the MC group of mice when compared to the CONTROL mice (Figure 2. F., p=0.028). Other beneficial resident gut bacteria including *Odoribacter splanchnicus*, a producer of short-chain fatty acids and bacterial sphingolipids in the intestinal microenvironment^72^, and *Turicibacter sanguinis* which produces intestinal serotonin^73^ were also found to be markedly decreased in relative abundance in the MC group when compared to the CONTROL group (Figure 2. G. and H., p=0.028). However, we also observed a significantly increased relative abundance of the beneficial gut resident *Bifidobacterium pseudolongum*^74^ in the MC-exposed mice compared to the CONTROL mice (Figure 2. D., p=0.0087). These results distinctly implied that early-life oral MC-administration in juvenile mice had a pronounced effect on the intestinal microflora in adulthood.

**Figure 2.**
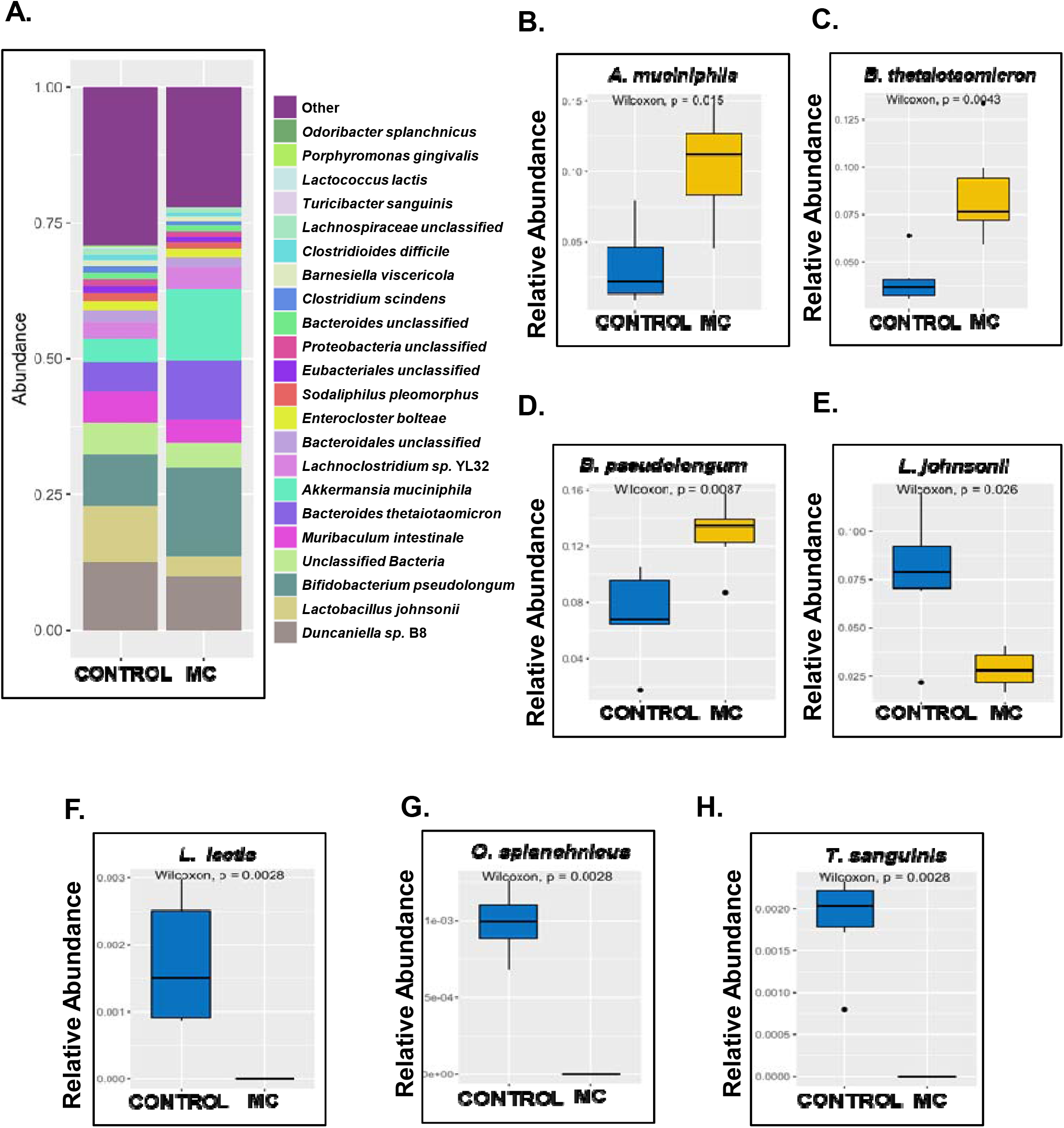
Relative abundance at the Species level. **(A.)** Bar plot showing relative abundance at the Species level for the two cohorts (CONTROL and MC). Box and whisker plots showing the differential abundance of **(B.)** *Akkermansia muciniphila*, **(C.)** *Bacteroides thetaiotaomicron*, **(D.)** *Bifidobacterium pseudolongum*, **(E.)** *Lactobacillus johnsonii* **(F.)** *Lactococcus lactis*, **(G.)** *Odoribacter splanchnicus*, and **(H.)** *Turicibacter* sanguinis.

### 3.2. Early life exposure to MC in mice leads to increased antibacterial resistance

After studying the effect of early exposure to MC in altering gut bacteriome, we hypothesized that MC may also influence the gut resistome. We observed that the alpha-diversity (Shannon index) of the gut resistome of mice in the MC group was significantly increased (p=0.0022) compared to the CONTROL group (Figure 3.A.). Tetracycline, aminoglycoside, macrolide, and glycopeptide were the most abundant resistant drug classes harboring maximum ARGs in both the mice groups (Figure 3.B.). The abundance of efflux pump encoding genes is a noted resistance mechanism used by several bacteria to expel toxic substances including antibiotics^75^. *bexA*, a multidrug efflux pump reported to confer resistance against fluoroquinolone antibiotics. Most of the Bacteroides species were significantly more abundant in the MC group than the CONTROL mice group^76^.

**Figure 3.**
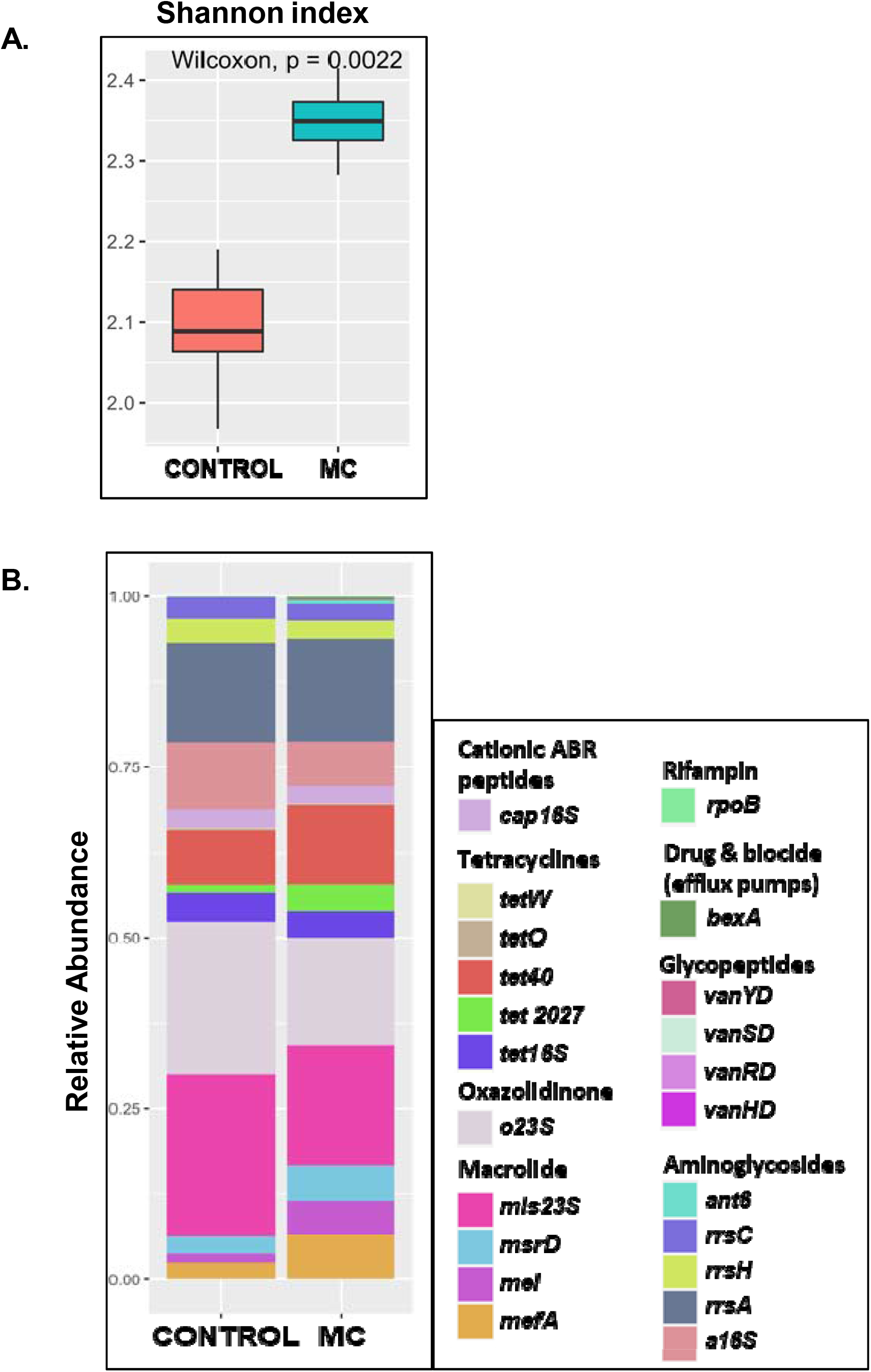
Diversity of Resistome. **(A.)** Microcystin-treated samples showed higher diversity in their resistome, both in terms of their evenness (Shannon index). **(B.)** Resistance gene abundance profile for CONTROL and MC cohorts.

Further, we studied the relative abundance of individual ARGs in the two experimental mice groups. *mefA* and *msrD* were significantly increased (p=0.0087 for *mefA* & p=0.015 for *msrD)* in the MC group (Figure 4.A. and B.). *mefA* and *msrD* are acquired resistance genes against macrolide antibiotics^77^. They are reported to be present in a large number of Gram-positive and Gram-negative bacteria^78^. Both ARGs are associated with the encoding of efflux pumps and are located together on MGEs^79–81^. *mel* which is an ARG against macrolides was also significantly increased (p=0.019) in the MC group compared to the CONTROL group (Figure 4.C.). Previous studies have reported that *mel* along with *mefA* and *msrD* and are located on the same MGE^82^. The relative abundance of ARGs, *ant6*, and *tet40* were also significantly increased (p=0.0043 for *ant6* & p=0.041 for *tet40)* in the MC mice group compared to CONTROL mice (Figure 4.D. and E.). *ant6* codes for resistance against aminoglycosides^83^. *tet40* confers resistance against tetracycline antibiotics and codes for efflux pumps. *tet40 is* also reported to be transferred among bacteria via HGT^84^. It is interesting to note that these mice were not exposed to antibiotics previously. Hence, our results from gut resistome analysis indicated that MC had a strong role in creating a selection pressure which resulted in increased antibacterial resistance which persisted for a long period. Resistance to above reported clinically important drug classes could lead to treatment failures in older ages in individuals exposed to MC at an early life.

**Figure 4.**
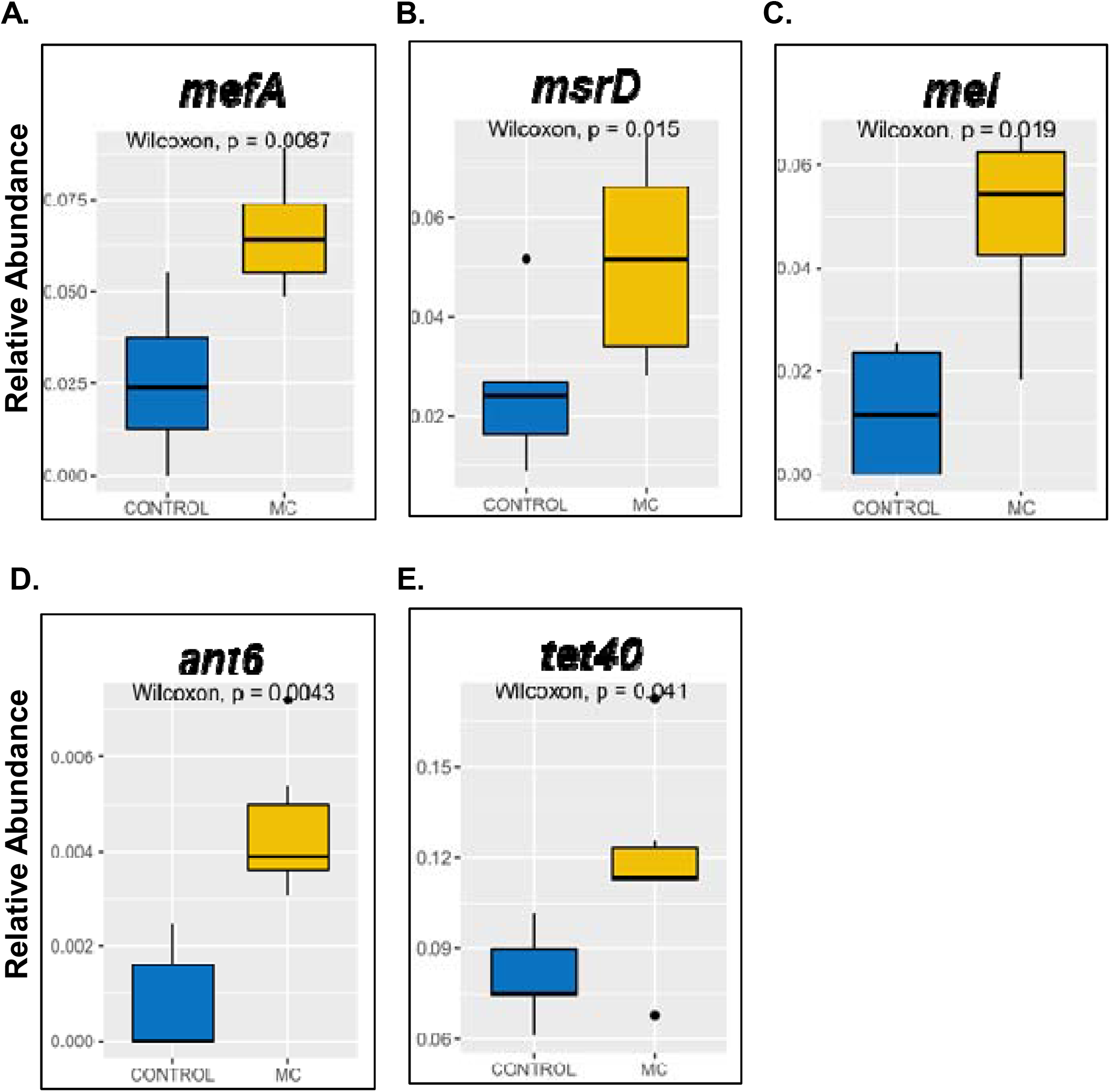
Box and whisker plots for differentially abundant resistance genes including macrolide-resistant class. [**(A.)** *mefA*, **(B.)** *mel*, and **(C.)** *msrD*], tetracycline-resistant class [**(D.)** *ant6*], and aminoglycoside-resistant class [**(E.)** *tet40*].

### 3.3. MC administration in mice decreases immunosurveillance but increases immunosenescence and systemic level of proinflammatory cytokine

Following MC-induced microbiome alteration and concomitant increase in ARG relative abundance, we wanted to observe the TLR-mediated intestinal immunosurveillance response as a result of early life MC exposure in the experimental groups of mice. qRT-PCR was performed using the intestine tissue samples from both WT CONTROL and MC groups. Our results showed significantly decreased gene expression of both TLR2 and TLR4 in the MC group when compared to the CONTROL group (Figure 5. A., ***p< 0.001). Interestingly, gene expression of the TLR-mediated anti-microbial peptide REG3G secreted by Paneth cells was also detected to be markedly decreased in the MC-exposed mice than CONTROL mice (Figure 5. A., ***p< 0.001). These results clearly indicated a weakened TLR-mediated immunosurveillance response in adulthood as a result of early life MC exposure in mice.

**Figure 5.**
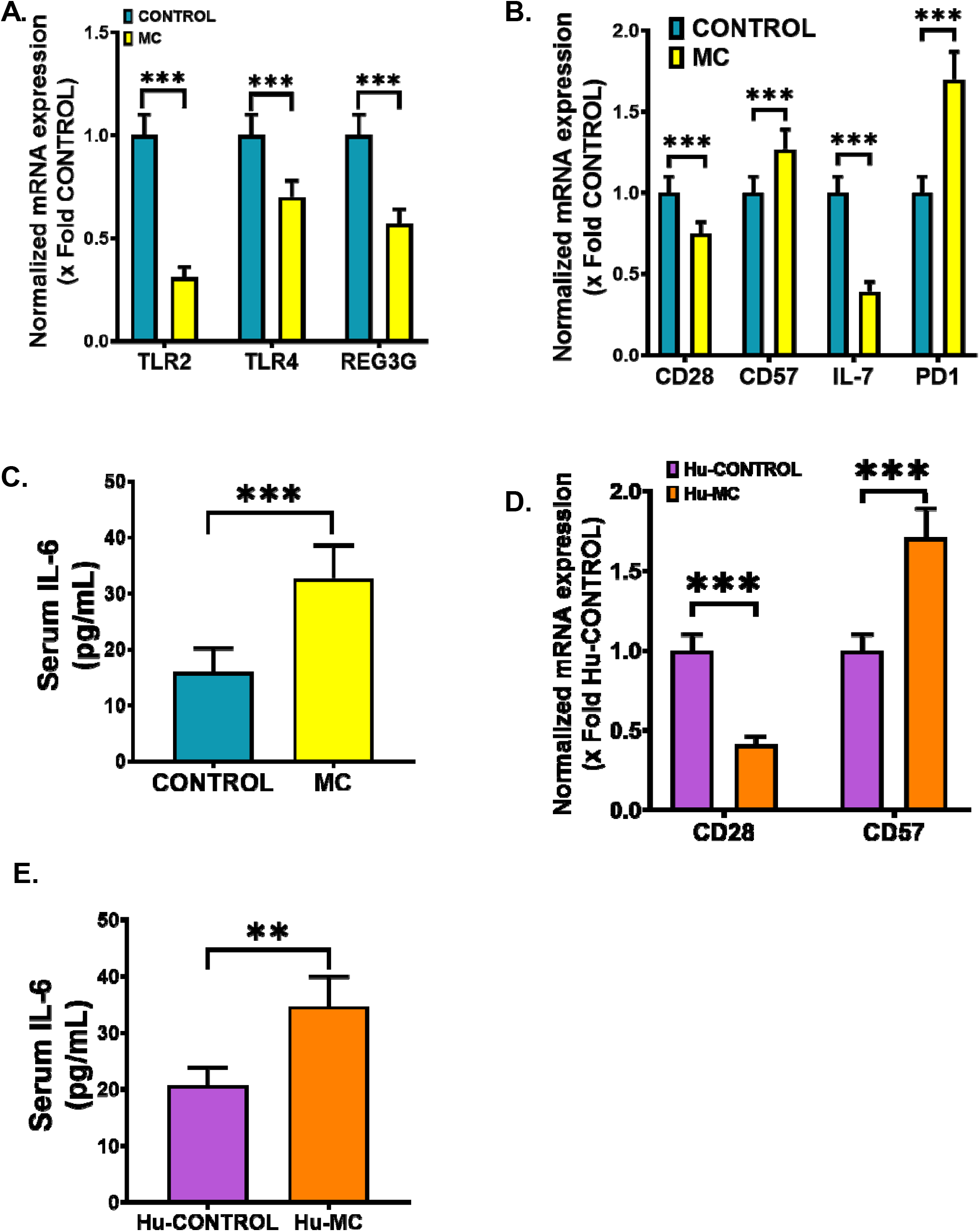
Early MC exposure caused pathological outcomes in the small intestine and elevated levels of systemic IL-6 in both WT mice and NSG^TM^ mice. Normalized mRNA expression of **(A.)** TLR2, TLR4, REG3G, and **(B.)** CD28, CD57, IL-7, PD1 against 18S in the small intestine of CONTROL and MC groups of mice and showed as the fold change of the CONTROL group (***p < 0.001). **(C.)** IL-6 (pg/mL) levels were measured in the serum of both CONTROL and MC mice groups and represented as bar graphs (****p< 0.001). Normalized mRNA expression of **(D.)** CD28, CD57 against 18S in the small intestine of Hu-CONTROL (NSG^TM^ mice treated with vehicle only) and Hu-MC (NSG^TM^ mice orally administered with MC for 2 weeks) groups of mice and showed as the fold change of the Hu-CONTROL group (***p < 0.001). **(E.)** IL-6 (pg/mL) levels were measured in the serum of both Hu-CONTROL and Hu-MC mice groups and represented as bar graphs (****p< 0.001). All data were represented as mean ± SEM, statistical significance was tested using unpaired t-test between the two groups, followed by Bonferroni Dunn Post hoc corrections.

Next, we wanted to study whether early life MC administration had any role in the intestinal T-cell immunosenescence process. Our qRT-PCR analysis showed a markedly decreased gene expression of both CD28 and IL-7 genes with a parallel elevated expression of CD57 and PD1 genes in the MC group when compared to the CONTROL group (Figure 5. B., ***p< 0.001). In addition, ELISA was performed using the serum samples of CONTROL and MC groups of mice to determine systemic IL-6 levels. Results indicated a significantly elevated (2.5-fold) circulatory IL-6 level in the MC group of mice compared to the CONTROL mice (Figure 5. C., ***p< 0.001). These results vividly implied that intestinal T-cell immunosenescence and systemic rise of the proinflammatory cytokine IL-6 in adulthood were promoted by early MC exposure.

To further validate the above-mentioned findings, we used NSG™ mice and MC-treatment was performed similarly to the WT mice to avoid any confounding result. As the NSG™ mice had hematopoietic stem cells engrafted in them, gene expression analysis of the intestinal T-cell immunosenescent markers was carried out by the qRT-PCR method using human primers. We detected a markedly lowered expression of the CD28 gene in the Hu-MC group of mice compared to the Hu-CONTROL group (Figure 5. D., ***p< 0.001) whereas CD57 gene expression was significantly elevated in the MC-exposed mice compared to the vehicle-treated CONTROL mice (Figure 5. D., ***p< 0.001). Circulatory IL-6 concentration was estimated by using a human-specific ELISA kit. Results showed a significantly elevated (1.67-fold) level of systemic IL-6 in the Hu-MC mice when compared to the Hu-CONTROL mice (Figure 5. E., ***p< 0.001). These results obtained from the NSG™ mice groups reflected exactly similar patterns and thus further confirmed the role of MC in modulating the intestinal immune system.

### 3.4. Association study between bacteria, ARGs, and immunological markers

To identify the probable source bacterium harboring the significantly increased ARGs we performed the gene provenance study. Results showed that *Bacteroides thetaiotaomicron* as the source bacterium harboring the macrolide resistance ARGs *mefA* (resistant to Erythromycin and Azithromycin)*, msrD* (resistant to Erythromycin, Azithromycin, and Telithromycin), and *mel* (resistant to Macrolides). This corroborates with our previous bacteriome analysis, where we found that *Bacteroides thetaiotaomicron* was significantly increased in the MC group. The aminoglycoside resistance gene *ant6* (resistant to Streptomycin) and tetracycline resistance gene *tetO* (resistant to Doxycycline, Tetracycline, and Minocycline) were found to be present in *Lachnospiraceae bacterium* A4, which was also found to be significantly increased in the MC group. Being gut commensals, the increased relative abundance of ARGs in these bacteria increases the risks associated with anti-microbial resistance due to its high transferability^85,86^.

Next, we wanted to study the association between MC exposure, ARGs, and altered immunological markers using the conditional independence test (Figure 6). We identified that both IL-7 and TLR2 were negatively affected by MC whereas ARGs *msrD, mefA*, and rpoB were positively affected by MC. *msrD, mefA*, and *rpoB* were negatively correlated to the expression of IL-7 (r=−0.736, p=0.006; r=−0.687, p=0.013; r=−0.584, p=0.045 respectively). Similarly, *mefA, ant6* and *rpoB* were negatively correlated to the expression of TLR2 (r=−0.743, p=0.005; r=−0.650, p=0.022; r=−0.617, p=0.032 respectively). MC was found to positively affect the expression of PD1, and serum IL-6 level as well as the ARGs *mefA, ant6, mel, tet40, bexA*, and *vanYD.* The anti-microbial resistant genes *mefA*, and *ant6* were found to be positively correlated to the expression of PD1 (r=0.662, p=0.018; r=0.686, p=0.013 respectively). Also, the ARGs *mel, tet40, bexA* and *vanYD* were found to be positively correlated to serum IL-6 level (r=0.760, p=0.004; r=0.662, p=0.018; r=0.821, p=0.001; r=0.625, p=0.029 respectively).

**Figure 6.**
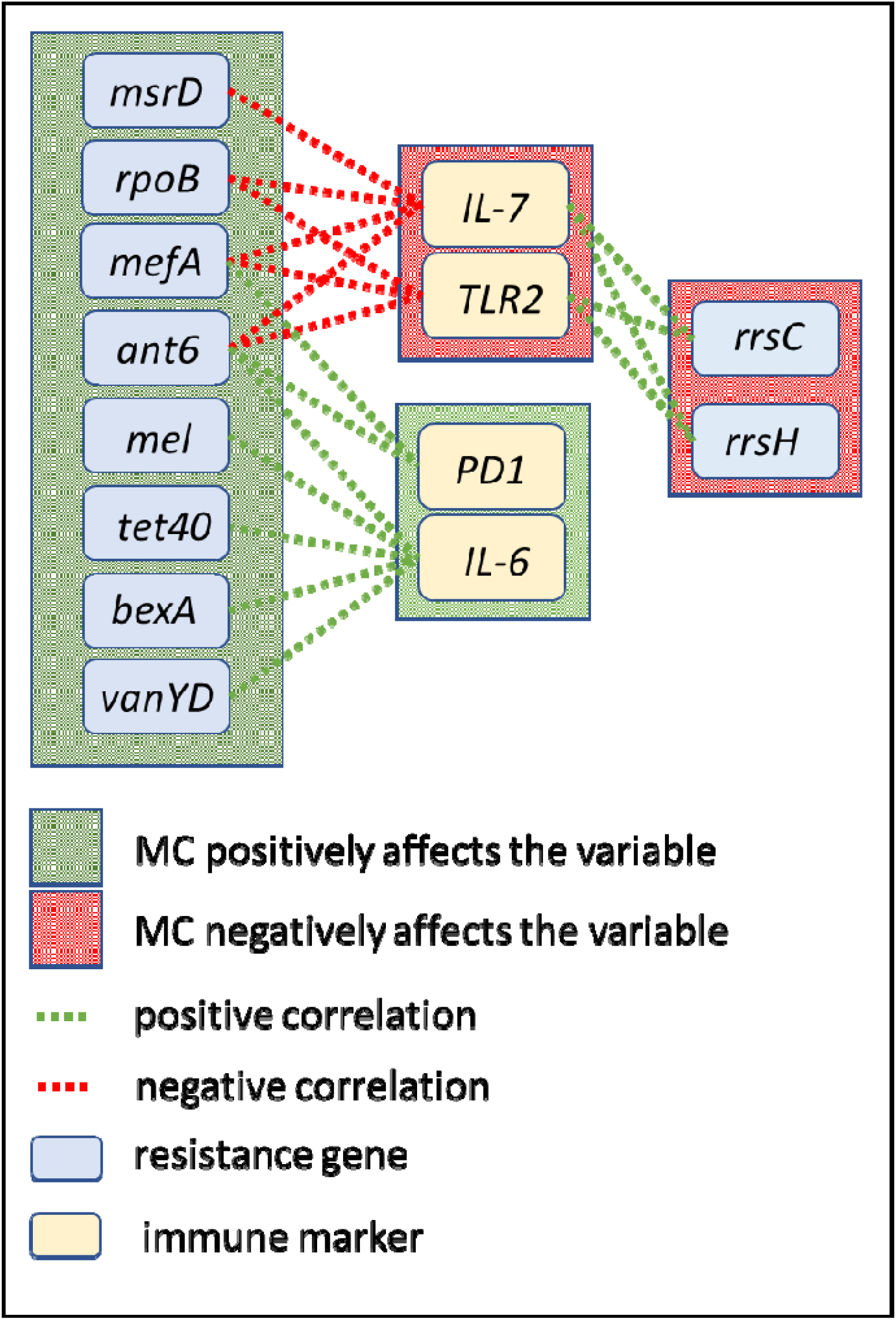
Correlated resistance genes and immune markers. Microcystin (MC) exposure changes the resistance profile of the gut microbiome and impacts the expression of immune markers. The abundance of antibiotic-resistant genes correlated strongly with that of immune markers. In the figure, green edges represent positive correlations and red represent negative correlations. However, that all the correlation values drop dramatically when conditioned on microcystin exposure. Conditional independence (CI) tests were performed with *bnlearn* R package with the Mutual Information method.

However, MC was found to negatively affect the gene expression of IL-7 and TLR2 as well as the ARGs *rrsC* and *rrsH.* Both *rrsC* and *rrsH* were found to be positively correlated to the expression of IL-7 (r=0.636, p=0.025; r=0.640, p=0.024 respectively) and TLR2 (r=0.614, p=0.033; r=0.636, p=0.025 respectively). These ARGs have been reported to be associated with *E. coli*, which were not detected in our bacteriome analysis^87,88^. These results showed that alteration of ARGs due to early life MC exposure has a strong effect on the expression of immunological markers related to systemic inflammation and immunosenescence, which corroborated with our previous qRT-PCR and ELISA results.

Further, we showed a co-occurrence network^89^ for both CONTROL (Figure 7.A.) and MC-treated (Figure 7.B.) samples. Correlations between microbes present in at least 50% of the samples (circles) and genes (squares) were shown. Microbial abundances and gene expressions were normalized separately, and we showed Pearson correlations (p=0.01) between these values [since these were separately normalized, they were not subjected to compositional anomalies^90^]. Green edges denoted positive correlations (meaning the microbial abundance increased with gene upregulation). Red edges denoted negative correlations (meaning microbial abundance increases with gene downregulation). Taxa were colored by phylum (Firmicutes-yellow, Bacteroidetes-dark purple, Proteobacteria-royal blue, Actinobacteria-brown, Verrucomicrobia-salmon). Networks were visualized using the Fruchterman-Reingold algorithm^91^, applied to the positive edges. We immediately noted the presence of a large Firmicutes cluster in both networks, double the size in the non-microcystin treated samples. In the CONTROL group, two genes (TLR2 and IL-7) were centrally located within the Firmicutes cluster, and a third (Serum IL-6) was tangentially located–indicating that as each of these immunological markers were up-regulated, the abundance of Firmicutes taxa within this large cluster was also increased. The smaller Firmicutes cluster in the MC-treated samples positively correlated with just one gene (TLR4). In both networks, these respective genes were negatively correlated with several Bacteroidetes taxa as well. In fact, in the CONTROL group, Firmicutes taxa were only the ones positively correlated with genes, and Serum IL-6 and Bacteroidetes were only negative. In the MC group, there were some exceptions (i.e., *F. rodentium* with TIM3, *B. thetaiotamicron* with PD1).

**Figure 7.**
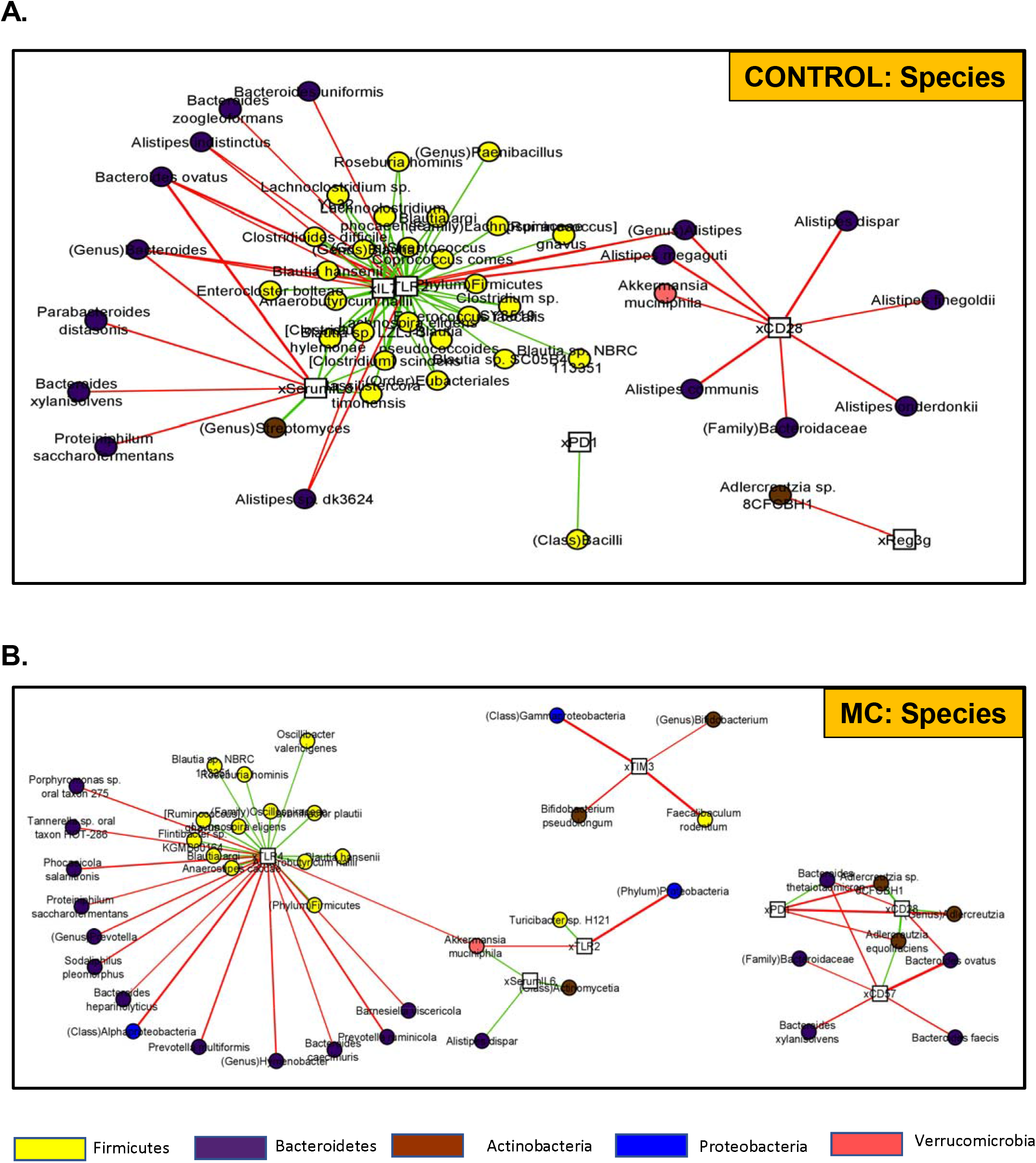
Correlation networks with the abundance of taxa and resistance genes. **(A.)** CONTROL and **(b)** MC groups co-occurrence networks built with Pearson correlations (p=0.01) between microbial abundances (circles) and gene expression values (squares), separately normalized to avoid compositional dependencies. Green edges indicate positive correlations, and red edges are negative correlations. Networks have been visualized with the Fruchterman-Reingold algorithm applied to positive edges.

Several other taxa exhibited differential behavior between the two networks. Multiple *Alistipes* taxa were negatively correlated with CD28 in the CONTROL samples, while in the MC group *Alistipes* was almost completely absent (only one taxon, compared to seven in the CONTROL group). In the MC group, we detected more involvement of *Actinobacteria*, particularly *Bifidobacterium* (negatively correlated with TIM3) and *Adlercruetzia* (negatively correlated with PD1). In contrast to its behavior in the CONTROL group, CD28 emerged as a gene that was upregulated with increased *Adlercruetzia* in the MC-treated group. CD57 also emerged in this network as negatively correlated with several Bacteroides taxa.

*Akkermansia muciniphilia* (Phylum Verrucomicrobia) was an interesting taxon, as it was reported as three times more abundant in the MC samples compared to the CONTROL group and its behavior was highly distinct between the two networks. In the CONTROL group, *A. muciniphilia* was simply another taxon negatively correlated with CD28 (along with several Bacteroidetes). In the MC group, *A. muciniphilia* actually increased with upregulation of Serum IL-6, and downregulation of TLR4 and TLR2.

In summary, dynamics involving Firmicutes and Bacteroidetes taxa with gene expression were similar in both networks, but much more polarized in the case of the CONTROL group. *A. muciniphilia, Alistipes, B. thetaiotamicron, Bifidobacterium*, and *Adlercruetzia* were all notable for their differential relationships with gene upregulation between the two networks. Specific genes involved in these relationships, and their behaviors, also varied greatly between the networks.

## 4. Discussion

We report the first-ever study of the effect of cyanobacterial bloom toxin on host gut resistome profiles and its subsequent association with possible dysregulated immune-phenotype. **Interestingly, the host resistome is increasingly becoming a target for deciphering the anti-microbial profile of subjects who are subjected to environmental exposures.** The resistome profile, the genes harbored within such host bacteria can be of immense consequence for the affected population who may be aging, have a compromised immune system, or may have susceptibility for gastrointestinal sepsis. The results reported in the study also have tremendous significance for clinics and hospitals where treatment of individuals with antibiotics in patients who have a compromised immune system or have an underlying disease with prior exposure to microcystin may have a poor prognosis.

The effect of environmental toxins on the gut microbiome has been of the growing interest of research^92^. Importantly, studies on the effect of environmental toxins in altering the gut microbiome and the associated health adversity are clinically significant not only to understand the pathology due to the cyanotoxin exposure but also in identifying future therapeutic targets. Cyanotoxins have a primordial position among the diverse environmental toxins studied due to the increasing exposures among humans and animals. Although MC-LR has prominent adverse effects on various organ systems including the liver, kidney, intestines, MC-LR exposure can also mediate significant changes to the host’s gut microbiome as observed by previous studies including our group^25,29,93,94^. We have shown previously that MC-LR treatment led to altered microbiome pattern^25^, resulting in increased lactate-producing bacteria in the gut microenvironment with elevated intestinal and serum lactate levels, which ultimately led to NADP Oxidase 2-dependent activation of TGFβ-mediated Smad2/3-Smad4 fibrotic pathway activation in the intestines of adult mice^26^. In the present study using wholegenome sequencing, we found that early life MC exposure leads to significant and persistent alteration of gut bacteriome and resistome in mice. The results clearly showed an association between altered resistome and bacterial species. Moreover, we have linked the increased ARGs due to early life MC exposure to the increased immunosenescence, systemic proinflammatory cytokine IL-6, and biomarkers related to innate immune response.

Early life MC exposure resulted in decreased (not significant) relative abundances of major phyla Bacteroidetes and Firmicutes. This is supported by the species level analysis where we observed that the relative abundances of *L.johnsonii, L.lactis*, and *T. sangunis* belonging to phylum Firmicutes and *O. splanchnicus* belonging to phylum Bacteroidetes were significantly decreased on MC exposure. These bacterial species have several beneficial functions on the host, hence a decrease in the abundance would be detrimental to the host gut health. Interestingly, we observed an increased relative abundance of *B. thetaiotamicron* also belonging to the Bacteroidetes phylum. *B. thetaiotamicron* is a known gut commensal that is primarily responsible for the carbohydrate metabolism of the host, converting complex sugars into simpler forms, thus producing various intermediate metabolites^95,96^. However, a study by Cutis et al.^85^ provided a mechanistic role of *B. thetaiotamicron* where the authors showed that fluctuation of sugar concentration in the intestines led to increased production of succinate by *B. thetaiotamicron*, which in turn augmented the pathogenicity of enteric pathogens. Relative abundance of *A. muciniphila* belonging to phylum Verrucomicrobia was significantly increased in the MC group compared to the CONTROL group. *A. muciniphila* has been reported to play an important role in maintaining tight junction integrity in the intestine and mucosal immune response as well as in host metabolism^97–99^ Studies have reported a decrease in the abundance of *A. muciniphila* during intestinal inflammation, especially in conditions like irritable bowel disease^100,101^. However, a separate study also reported that a decrease in gut mucus barrier strongly correlated with increased abundance of *A. muciniphila* which increased the susceptibility to colonization by pathogenic bacteria^102^. Hence, we need an in-depth mechanistic study to understand the exact roles of *B. thetaiotamicron* and *A. muciniphila* in MC exposure in the future. The species-level analysis also showed that the relative abundance of *B. pseudolongum* of Actinobacteria phylum was significantly increased in the MC group. *B. pseudolongum*, is a known probiotic having an anti-inflammatory effect and mediates immune homeostasis^103,104^. An increase in abundance of *B. pseudolongum* could possibly be an adaptive response by the host in order to protect the intestinal microenvironment from the damages due to MC exposure. A limitation in our study to fully understand some of our differential abundance of the above-mentioned species might be the sample size and can be perceived as a limitation in this study.

An altered host gut resistome is associated with gut bacterial alterations which increase the chances of developing antibiotic resistance. The important highlight of this study is the increase in antibiotic resistance due to early life exposure to MC. There is little or no evidence in the existing literature on MC exposure in altering host gut resistome. Our results clearly showed that MC exposure significantly increased the α-diversity of ARGs and these changes had a prolonged effect pertaining to the observed results even after 10 weeks. Increased resistance to multiple drugs like tetracycline, aminoglycosides, glycopeptides, and macrolides which are used to treat both Gram-positive and Gram-negative bacterial infections in general clinical settings^105^ due to early life MC exposure is of serious concern. We would also like emphasize that the mice used in these study were raised in a controlled environment without any prior antibiotic exposures, hence the change in ARGs and the effects of the change in resistome can be ascribed solely to the exposure of the environmental HAB toxin.

Individual ARGs that were significantly increased due to MC exposure had high transferability. ARGs can be transferred between bacteria by HGT using MGEs like transposons, integrons, or via plasmids and chromosomes ^38,40,106^ According to previous studies, *mefA, msrD*, and *mel* coding for the macrolide resistance were reported to be present in the same MGE transposon. Similarly, *ant6*, an ARG resistant against aminoglycosides is present on transposons, plasmids, and chromosomes^107^. The ARG *tet40* codes for tetracycline resistance were reported to be present in transposons and plasmids^108^. These ARGs possess the risk of spreading resistance against these antibiotics which could be lethal to the host health in the future rendering the host to be untreatable by these broad-spectrum antibiotics in future infections.

Our bacteriome and ARG analysis also suggested that MC exposure may be creating a potential selection pressure allowing certain species with the advantage of ARGs to survive. Gut commensals are reported to possess ARGs, which could be transferred by HGT^109^. In our study, *B.thetaiotamicron*, which was significantly increased on MC exposure, was found to harbor *mefA, msrD* and *mel* genes by gene provenance study. Studies suggested that bacterial species belonging to phylum *Bacteroidetes* are able to accumulate ARGs and transfer them to other gut commensals and pathogens^110^. Similarly, *ant6* and *tetO* were found to be associated with *Lachnospiraceae bacterium A4.* This result is of immense importance as it clearly shows that exposure to cyanobacterial toxins like MC increases the likelihood of spreading ARGs among neighboring commensals, opportunistic pathogens, or even to enteric pathogens during future bacterial infections like *Vibrio sp.* increasing the antibiotic resistance in the gut microbiome.

Next, we wanted to link the altered microbiome and resistome signature to the host’s TLR-mediated immunosenescence, innate immune response, and systemic inflammation. Immunosenescence is part of the normal physiological process where adaptive immunological response grew weaker with age-related changes in the body^111^. Importantly, a study unveiled that the continuation of a healthy gut microflora along with consumption of pre and probiotics might be effective in delaying aging-related immunosenescence and inflammation^112^. CD28 is a co-stimulatory receptor on present on T-cells which plays a major role in naïve T-cell activation^113^ whereas CD57 is another T-cell receptor that significantly increased in expression during later stages of life^114^. Both CD28 and CD57 serve as extensive hallmarks of immunosenescence as increased CD8^+^ CD28^-^ CD57^+^ T-cell populations are identified and reported in various age-related pathological conditions^115–117^. Similar results were also detected in our study as we observed increased gene expression of CD57 with a parallel decreased gene expression of CD28 in both MC-exposed WT and humanized NSG^TM^ mice. Additionally, IL-7 and PD1 can also serve as considerable markers of immunosenescence. IL-7 and its receptor-mediated signaling is a key growth factor needed for the T-cell development process^118^ that decreases with age, whereas the PD1 receptor acts as a major checkpoint for controlled T-cell mediated immune response^119^, and its expression eventually increase in the elderly population^120^. Similar decreased expression of IL-7 with increased PD1 gene expression as obtained in our study in combination with the previously mentioned CD28, and CD57 gene expression results clearly indicate the MC-exposure during childhood and resulting dysbiosis can potentiate early onset of immunosenescence even at a relatively young age. Toll-like receptors are one of the primary components of innate immune response that significantly contribute to the host’s immunosurveillance mechanisms^121^. However, gene expressions of both TLR2 and TLR4 were decreased in our study as a result of early life MC exposure, which suggests an impaired immunosurveillance mechanism in the intestinal microenvironment. In turn, this MC-mediated impaired immunosurveillance possibly led to decreased gene expression of the anti-microbial peptide gene REG3G, which is a TLR signaling-dependent process^122^. In addition, we also observed an elevated level of circulatory IL-6 in both MC-exposed juvenile WT and adult NSG^TM^ mice, indicating that this systemic inflammation exerted by MC administration was completely independent of the age of mice. Increased systemic level of IL-6 level has been associated with various infectious and non-infectious disease pathologies^123,124^ and we have already reported it as an extensive soluble mediator of gut-brain axis dependent neuroinflammation^44,125^. However, an in-depth mechanistic study demonstrating the role of MC-driven IL-6 trans-signaling and resulting inflammatory surge connecting gut-brain axis might shed further light in the field of cyanotoxin research.

Further, we have identified a direct association between the altered biomarker and the ARGs due to early life MC exposure. MC exposure directly influenced the expression of PD1, which further increases the chances of immunosenescence in later life. Expression of PD1 had a positive correlation with ARGs *mefA* and *ant6.* Expression of systemic IL-6, a pleiotropic cytokine was found to be directly influenced by MC and 5 ARGs. This strongly suggests that increased genes resistant to multiple antibiotics may have a direct influence on the proinflammatory phenotype due to MC exposure. Results from this association study also showed that expression of IL-7, a negative regulator of immunosenescence decreased with the increase of certain ARGs. TLR2, a marker of innate immune response and immunosurveillance, was also found to be decreased with the increase of specific ARGs. Apart from the association between ARGs and immunological markers, we have also reported the association between altered gut bacteriome as a result of MC treatment and host immunological markers. This is the first time that an association with the expression of biomarkers and ARGs are being reported in an MC exposure study. However, more mechanistic studies need to be conducted to further confirm the association between the two factors.

In conclusion, our study reveals a new aspect in MC exposure pathology whereby MC exposure in early life poses a high risk of developing antibiotic resistance which persists in the later adult life in the experimental mice. Incidences of co-infection by potentially pathogenic bacteria like *Vibrio vulnficus* and *Vibrio parahaemolyticus* has been reported to occur in human after exposure to cyanotoxins like microcystin^126–128^. This led to higher rates of mortality, especially in cases where the exposed individuals were severe immunocompromised ones^129–131^. Our study clearly showed that MC exposure in early life increased multi-drug resistance to clinically important antibiotics which are also used in the treatment of *Vibrio* infection like tetracycline^132^. It increases immune dysfunction in adult stages as seen by an increase in the expression of immunosenescence markers at the end of the experimental time period, which in turn are directly associated with the increase in antibiotic resistance. Hence, this study provides novel insights for identifying therapeutic targets which can be utilized in the future in treating pathological conditions due to the altered gut bacteriome during MC exposure as well as in co-infection like vibriosis.

## Funding

This study was supported by NIH grant 5P01ES028942-04, Subproject-5262, to Saurabh Chatterjee (Project PI).

## Author contributions

S.C., P.S., D.B. conceived and designed research. P.S., D.B., R.S. V.S., performed the experiments. P.S., D.B., S.C., G.S., K.M, T.C., analyzed the data and interpreted the results of experiments. P.S., V.S., and D.B. prepared the figures. P.S., D.B, and S.C. drafted the manuscript. G.I.S, D.E.P., and B.B. edited the manuscript. S.C. edited, revised, and approved the final version of the manuscript. The authors read and approved the final manuscript.

## Acknowledgment

The authors gratefully acknowledge the technical services at CosmosID Inc. (Germantown, MD, USA) for microbiome sequencing, and AML Labs (St. Augustine, FL, USA). We also thank the Instrumentation resource facility (IRF) at the University of South Carolina for equipment usage and consulting services.

## Declaration of competing interest

The authors declare that there is no conflict of financial or competing interests.

## Notes

### Competing Interest Statement

The authors have declared no competing interest.

